# Learning to quantify uncertainty in off-target activity for CRISPR guide RNAs

**DOI:** 10.1101/2023.06.02.543468

**Authors:** Furkan Özden, Peter Minary

## Abstract

CRISPR-based genome editing technologies have revolutionised the field of molecular biology, offering unprecedented opportunities for precise genetic manipulation. However, off-target effects remain a significant challenge, potentially leading to unintended consequences and limiting the applicability of CRISPR-based genome editing technologies in clinical settings. Current literature predominantly focuses on point predictions for off-target activity, which may not fully capture the range of possible outcomes and associated risks. Here, we present crispAI, a neural network architecture-based approach for predicting uncertainty estimates for off-target cleavage activity, providing a more comprehensive risk assessment and facilitating improved decision-making in single guide RNA (sgRNA) design. Our approach makes use of the count noise model Zero Inflated Negative Binomial (ZINB) to model the uncertainty in the off-target cleavage activity data. In addition, we present the first-of-its-kind genome-wide sgRNA efficiency score, crispAI-aggregate, enabling prioritization among sgRNAs with similar point aggregate predictions by providing richer information compared to existing aggregate scores. We show that uncertainty estimates of our approach are calibrated and its predictive performance is superior to state-of-the-art *in silico* off-target cleavage activity prediction methods.

## Introduction

CRISPR/Cas9 system (Clustered regularly interspaced short palindromic repeats/ CRISPR-associated protein 9) first discovered in the immune mechanisms of bacterial and archeal species [1] and quickly became among the most popular gene editing technologies recently with successful applications of editing eukaryotic genomes [2].

CRISPR/Cas9 system has also been applied to knockout-screening studies [3], accelerating the understanding of variant and gene functions by uncovering causal relations between mutations and phenotypes. Due to its high efficiency, simpler design and easier operation procedures in comparison to earlier genome editing methods like Zinc Finger Nucleases (ZFNs) and Transcription Activator-Like Effector Nucleases (TALENs), it is becoming a standard in genome engineering field and has the potential to lead new treatments for genetic diseases [4].

The CRISPR/Cas9 system works by using a programmed single guide RNA (sgRNA) to direct the Cas9 enzyme to a specific target sequence in the targeted DNA site. Once the Cas9 enzyme is bound to the target DNA, it creates a double-strand break, which triggers natural repair mechanisms of the cell. This can result in the targeted sequence being edited by deletion, insertion, or replacement, depending on the desired outcome. However while CRISPR/Cas9 system operates on the targeted DNA region, cleavage may also occur at other genomic loci with a DNA that is not fully complementary to the sgRNA and has several base mismatched sites with the sgRNA. These cleavage effects, referred to as ‘off-target’ cleavage, are unintended and can be dangerous, resulting in unintended changes in the genome, leading to unwanted gene mutations and potentially harmful effects [5]. The existence of the off-target cleavage phenomenon has been one of the key factors to limit the development and applicability aspects of the CRISPR-based genome editing systems. Studies found out that few-mismatch DNA sites are potentially recognizable by the sgRNA during the guiding process [6]. Also, the off-target effects have been shown to be dependent on many other factors such as nucleosome occupancy, chromatin accessibility and both binding and heteroduplex energy parameters of the sgRNA of choice [7]. However, there are many potential sgRNA sequences that can direct the system for the intended cleavage effect to take place and hence one of the key design aspects of CRISPR-based systems is to evaluate and assess the error profile of the sgRNA of interest.

Up to date, many off-target cleavage activity prediction tools have been proposed to predict the potential off-target activity of a given sgRNA-target pair (i.e., targeted DNA site and corresponding single guide RNA sequences). These algorithms use the data generated by experimental CRISPR off-target detection assays, such as GUIDE-seq [8], CHANGE-seq [9], DIGENOME-seq [10], CIRCLE-seq [11], and predict a point score for off-target cleavage activity for a given sgRNA-target pair based on the training data. These tools can be divided into two main categories: (i) Conventional Machine Learning-based models and (ii) Deep Learning-based models. Conventional machine learning models have been extensively used for on-and off-target prediction in CRISPR/Cas9. Various algorithms such as random forest, SVM [12], logistic regression [13], gradient boosting [14], and ensemble learning [15] have been employed to predict off-target activity. While conventional machine learning models have shown promising results, recent studies using deep learning techniques have demonstrated even better performance [16]. These models utilize novel sequence encoding strategies, feature engineering approaches by introducing physical features [17], class rebalancing techniques, and attention mechanisms [18] to improve prediction performance. While several models have been developed, most of them are primarily dedicated to the classification task, aiming to predict the activity status of the sgRNA-target interface. Notably, MOFF score [19] stands out as a top-performing model which tackles the regression task of predicting the activity level of the sgRNA-target interface, which is the focus in this article as well. For a more detailed review of the current literature Sherkatghanad et al. [20] presented a detailed overview of machine learning and deep learning-based studies for on/off-target activity prediction task for CRISPR/Cas9 systems.

It is worth noting that, the imbalance in CRISPR off-target prediction data poses a significant challenge, as the number of true off-target sites recognized by whole-genome detection techniques is much smaller than that of all possible nucleotide mismatch loci [20, 21]. This imbalance can make training routine machine learning models difficult, resulting in high accuracies for the majority class but poor performance for the minority class, which is of greater interest in this context because it represents the actual off-target sites that can lead to unintended consequences in genome editing [14]. These point predictions, while informative, may not fully capture the range of possible outcomes or the associated risks in the editing process. To the best of our knowledge, the only other study that incorporates uncertainty estimates into the off-target cleavage activity prediction task is Kirillov et al. [22], where authors trained a Gaussian Process Regression model. Incorporating uncertainty estimates into predictive models would facilitate the identification and prioritization of potential off-target sites, especially when they have similar point off-target activity predictions. By considering the uncertainty estimates, researchers and practitioners can differentiate between sites with similar point predictions and prioritize those with lower uncertainty, thereby reducing the chances of unforeseen off-target effects. This prioritization strategy, enabled by uncertainty estimates, would lead to improved validation and optimized guide RNA design, reducing potential risks associated with CRISPR-based genome editing applications. Additionally, genome-wide off-target detection methods encounter various experiment-specific limitations that can affect their sensitivity. One of the key contributing factors responsible from the highly imbalanced nature of the off-target data produced by such detection techniques stems from the fact that these methods rely on Next Generation Sequencing (NGS)-based DNA sequencing assays which generally exhibit high sparsity, often encountered in microbiome, bulk-, and single-cell RNA experiments [23, 24]. These limitations hinder the ability to differentiate between real biologically inactive off-target sites and technical errors, resulting in false negatives in the analysis for the guide RNA of interest [25]. Addressing the highly imbalanced nature, and the uncertainty, of the off-target cleavage data is crucial, particularly because mistaking an active off-target site for an inactive one can have significant consequences. Such errors can disrupt cellular function or confound experimental interpretation, whereas mistaking an inactive site for an active one may only necessitate designing another gRNA [14]. In this work we present crispAI, to accurately predict the off-target cleavage activity in a probabilistic framework, allowing for quantification of the uncertainty in the predictions and crispAI-aggregate to provide a probabilistic genome-wide specificity estimate for a given sgRNA.

## Results

### Overview of crispAI

We designed crispAI to be a hybrid deep learning architecture based on the count distribution Zero-Inflated Negative Binomial (ZINB), which accounts for highly imbalanced nature of the off-target cleavage data and models the off-target cleavage activity in a probabilistic framework (Methods). Our architecture uses a combination of Convolutional Neural Network (CNN) and bi-directional Long Short Term Memory (biLSTM) layers to extract sequence-based features of the sgRNA-target pair, which are encoded using a binary matrix encoding scheme first presented by Lin et al. [26]. We used an additional CNN layer to process physical descriptors of the sequence context, namely: (i) Block Decomposition Method (BDM) score [27]; (ii) GC content; (iii) NuPoP Occupancy; and (iv) NuPoP Affinity scores [28]. The importance of these descriptors for off-target cleavage activity has been highlighted by a recent study [7]. Both sequence based features and physical descriptor features are used to extract features related to the off-target cleavage activity of the sgRNA-target interface. The extracted features are then concatenated in a late fusion fashion for the final Fully Connected (FC) layer to predict three parameters, *µ, θ* and *σ*, associated with the ZINB distribution. We trained the network weights based on a loss function optimising the likelihood of the observed data. (Fig. 1).

**Fig. 1.**
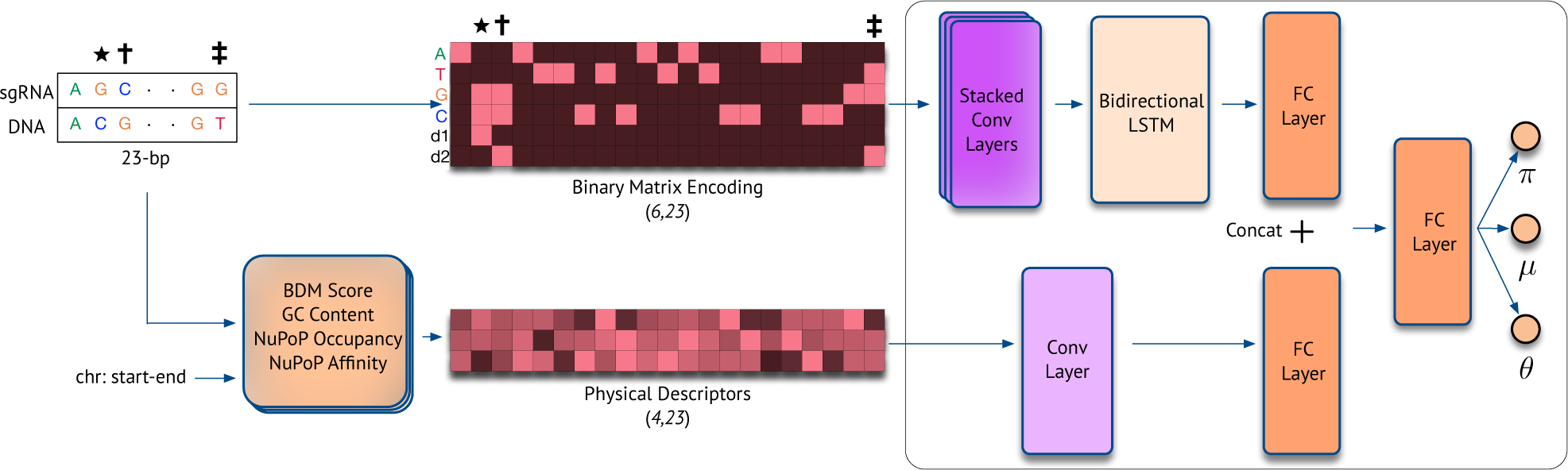
Workflow Figure. Logical OR operation is employed to encode the sgRNA-target interface to a matrix of shape (6, 23) where the first 4 rows represent sequence letters (i.e., ATGC) and the last 2 rows represent mismatch direction. Physical descriptors of the sequence context such as: (i) BDM Score; (ii) GC content; (iii) NuPoP Occupancy; and (iv) NuPoP Affinity are calculated and normalized values are encoded into a matrix of shape (4, 23) for each sequence position. Then, the sequence encoding features are extracted with a series of Convolutional (Conv) layers followed by a Bi-LSTM layer and features of physical descriptors are extracted with a Conv layer. Both extracted feature vectors are mapped to 128 *− d* vector spaces with Fully Connected (FC) layers and are concatenated. The concatenated features are then passed to a final FC layer to predict three parameters *π*, *µ* and *θ*.

### Enabling uncertainty quantification for off-target activity prediction

Our proposed architecture, crispAI, models the off-target cleavage activity in a probabilistic framework. Hence, making it possible to sample from the posterior off-target activity distributions conditioned on both the sequence-based and physical features of the sgRNA-target interface. Fig. 2 depicts example distributions for eight randomly selected test samples from the left-out test portion of the CHANGE-seq [9] dataset, where point predictions and the ground truth CHANGE-seq detected off-target cleavage activity values are shown with vertical lines.

**Fig. 2.**
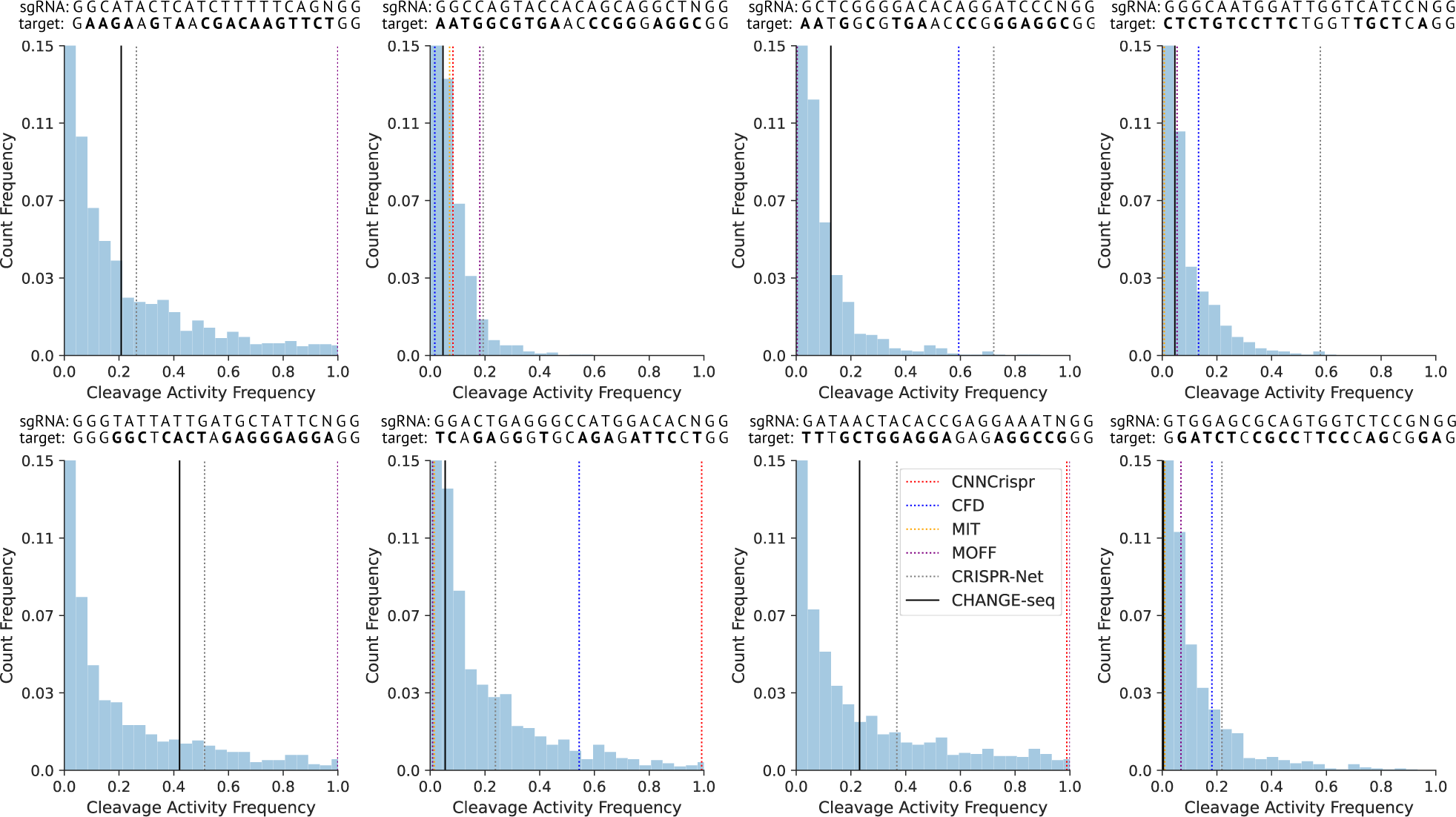
Example distributions for eight randomly selected sgRNA-target interfaces. Our approach, crispAI, enables sampling from the posterior off-target activity distribution conditioned on the sequence-based features of the sgRNA-target pair of interest. Vertical lines represent CnnCrispr score, CFD score, MIT score, MOFF score, CRISPR-Net score and ground truth CHANGE-seq detected off-target cleavage activity frequencies.

We started evaluating crispAI by visualising the uncertainty estimates for individual sgRNA-target pairs on the left-out test portion of the CHANGE-seq dataset [9]. CHANGE-seq is a highly scalable, Next Generation Sequencing (NGS)-based *in vitro* genome-wide Cas9 off-target detection assay which includes both epigenetic and genetic impact. Authors identified 202, 043 off-target sites for 110 sgRNAs on 13 therapeutically relevant loci in human primary T-cells. We randomly splitted 10% of the CHANGE-seq dataset for testing obtaining 168, 465 samples in total. First, we obtained posterior parameters of the respective ZINB distributions conditioned on the sgRNA-target pair features for each sample using crispAI on the left-out test portion of the CHANGE-seq dataset. We sampled corresponding posterior distributions many times to obtain empirical Probability Mass Functions (PMFs) for each sample. Then, we obtained CnnCrispr score [16], CFD score [12], MIT score [29], MOFF score [19], CRISPR-Net score [26] on the same test portion for comparison with the ground truth CHANGE-seq detected activity values and expected value of the crispAI-predicted PMFs. We sorted the test samples and all associated point predictions along with the ground truth values based on their predicted Upper Confidence Bound values (UCB). Fig. 3**a**. shows the confidence intervals of crispAI-predicted posterior PMFs, point predictions of competing methods and the expected value of the predicted PMFs and the ground truth CHANGE-seq detected activity values. We observed that predicted intervals accurately captured the ground truth cleavage activity values for almost all of the samples. For on-target sites and highly active off-target sites (e.g., activity frequency *>* 0.25) the posterior distributions yielded very high 95% CI UCBs reaching up-to maximum activity frequency, which is expected since the detected frequency value highly depends on the number of other detected off-target sites for the same guide RNA instead of the individual features of the interface itself for highly active off-target sites and on-target sites [9]. Additionally, we observed that the expected values of crispAI-predicted PMFs closely resembled the ground truth activity values, whereas CnnCrispr score, CFD score, MIT score and CRISPR-Net scores are generally far off while MOFF score is somewhat better at distinguishing between minimal activity and highly active sites. Due to vast imbalance between minimal activity off-target sites (e.g., *<* 0.007) and higher activity off-target sites we splitted the comparison into two parts. Fig.3**b.** shows the samples with ground truth CHANGE-seq detected activity values *>* 0.007 for better visibility.

**Fig. 3.**
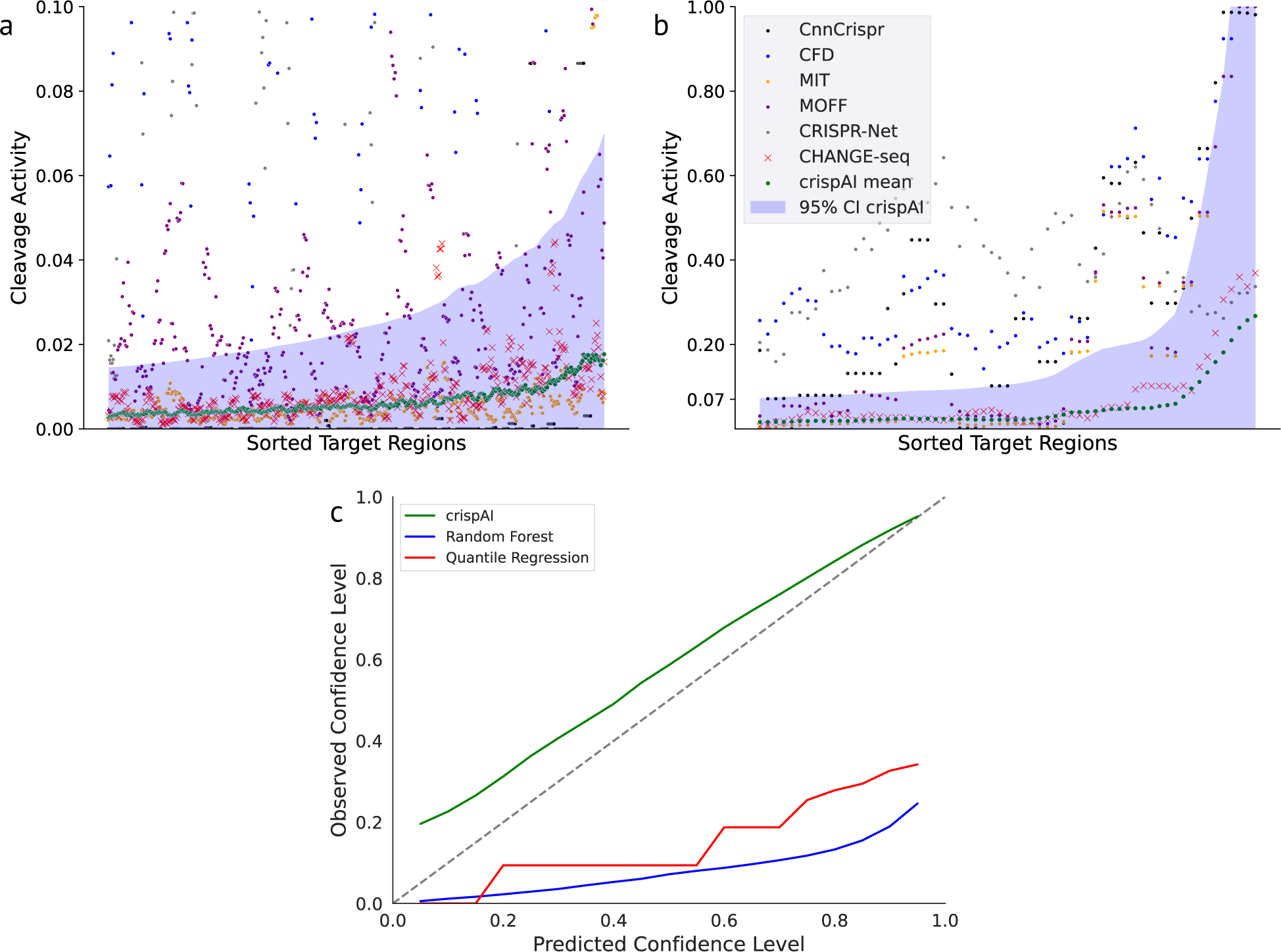
Depiction of crispAI-predicted uncertainty estimates on the test portion of the CHANGE-seq data. **a.** First, we obtained the crispAI-predicted off-target activity distributions for samples with positive CHANGE-seq counts. Then we calculated both the 95% confidence interval and the expected value (green) for each distribution and sorted the off-target samples and associated predictions of CnnCrispr score (black), CFD score (blue), MIT score (red), MOFF score (purple), CRISPR-Net score (gray) along with the ground truth CHANGE-seq (red cross) detected activity (normalized between 0 and 1) in the increasing order, based-on the predicted Upper Confidence Bound (UCB) values. Due to vast imbalance of low-activity samples (e.g., y *<* 0.07) the confidence interval plot is splitted to two parts. **b.**Part **a.** is repeated for cleavage activity values greater than 0.07. **c.** Observed vs. Predicted confidence level plot is depicted as the uncertainty diagnostics plot. For benchmarking, we generated baseline uncertainty estimates by training Random Forest and Quantile Regression models on the same training portion of the CHANGE-seq dataset.

Ideally, in single guide RNA design, a maximum cleavage activity on the targeted site with minimum activity on off-target sites is desired. Hence, Upper Confidence Bounds (UCB) of uncertainty estimates for the prediction of on-target activity task are not of concern and should be set to maximum activity value possible, measuring the Lower Confidence Bound (LCB) accordingly for the desired Confidence Interval (CI). Similarly, for LCB of uncertainty estimates for the prediction of off-target activity task are not of concern and should be set to lowest activity value possible - again, measuring the UCB for off-targets accordingly for the desired CI. Therefore, we set LCBs of all off-target samples at minimum activity (i.e., 0) and we measured the UCB at 95% CI.

Next, to evaluate the quality of the uncertainty estimates of crispAI using the same test portion of the CHANGE-seq dataset, we computed the diagnostic calibration measure for uncertainty estimates proposed by Kuleshov et al. [30]. Proposed diagnostic measure suggests that a well-calibrated uncertainty forecast should contain *N−*percent of samples in *N−*percent confidence interval. By plotting the observed confidence level against the expected confidence level, we produced the proposed diagnostic plot, in which well-calibrated uncertainty estimates are expected to produce a straight line. For comparison with baseline uncertainty estimates, we trained Random Forest regressor and Quantile Regression models on the same data crispAI is trained. Figure 3**c**. depicts diagnostic lines for each method. We observed that diagnostic plot of crispAI-predicted CI levels is similar to the ideal calibration line, while baseline uncertainty estimates are far off.

### Improving *in silico* CRISPR/Cas9 off-target cleavage activity prediction performance with crispAI

To evaluate the predictive performance of crispAI, we used 5 test datasets: (i) left-out test portion of the CHANGE-seq dataset [9]; (ii) GUIDE-seq dataset [8]; (iii) SITE-seq dataset [31]; and the datasets presented in Chuai et al. [32] (iv) HEK293T cell-line; and (v) K562 cell-line (Datasets). CHANGE-seq is a scalable, automatable tagmentation-based method for measuring the genome-wide activity of Cas9 in vitro. GUIDE-seq is a method for globally detecting DNA double-stranded breaks introduced by CRISPR RNA-guided nucleases (RGNs), using the capture of double-stranded oligodeoxynucleotides to identify off-target cleavage activities. SITE-Seq is a biochemical method for identifying off-target cleavage sites of CRISPR-Cas9 RNA-guided endonucleases through the selective enrichment and sequencing of adapter-tagged DNA ends. We inputted sgRNA sequence, target sequence and associated coordinates of each sgRNA-target pair in the mentioned datasets to crispAI pipeline, and obtained crispAI-predicted posterior off-target activity score distributions as depicted in Fig.4**a**.

We observed that point predictions obtained using the expected values of crispAI-predicted distributions, significantly out-performed all of the competing tools with respect to Spearman correlation coefficient on all test datasets except SITE-seq and performed second-best following CnnCrispr for this dataset. Specifically we observed, 21.09%, 6.92% and 17.77% improvements over the best models in HEK293T, K562 and union of the two test datasets respectively (Fig.4**b**). Similarly for the CHANGE-seq and the GUIDE-seq datasets we observed, 19.51%, 10.76% improvements over best performing tools MOFF and CRISPR-Net respectively and 12.01% deterioration on SITE-seq dataset performing second best behind best performing tool CnnCrispr. Additionally, we plotted 5 box-plots, one for each dataset, illustrating the distribution of Coefficient of Variation of crispAI-predicted distributions to compare with the variances of the ground truth cleavage activity values given in the datasets. We observed increased median lines and 3rd quartile lines as the variance of the dataset increases as expected.

**Fig. 4.**
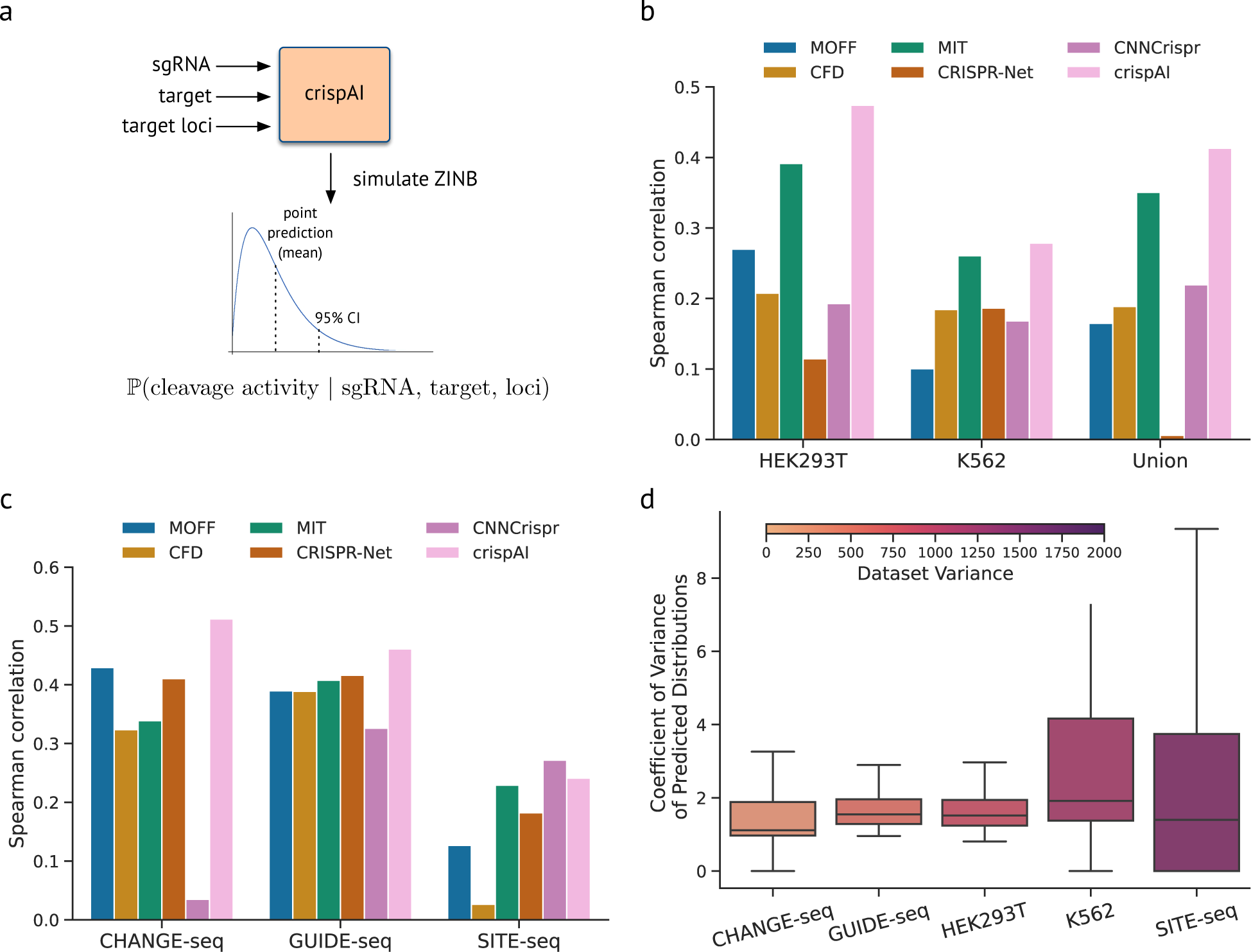
Off-target cleavage effect prediction with crispAI. **a.** Input/Output schematic representation of crispAI. 23*−*bp sgRNA and target DNA sequences with target DNA coordinates of the interface are inputs to our model. The model is trained to predict ZINB associated posterior parameters conditioned on the inputs. The conditional posterior distribution is then sampled. **b.** Predictive performance comparison of competing models on HEK293T, K562 cell-line datasets and their unions comprising of *n* = 536, 120 and 656 samples respectively. Bar-plot displaying Spearman correlation coefficient between the ground truth cleavage values in the respective datasets and the predictions of MOFF score, CFD score, MIT score, CRISPR-Net score, CNNCrispr score and crispAI score. For performance comparison, expected value of crispAI-predicted posterior distributions are used as point predictions. **c.** Similarly to part **b.** performance of aforementioned methods are compared on CHANGE-seq, GUIDE-seq and SITE-seq datasets comprising of *n* = 168, 465, 443 and 6, 097 samples respectively. **d.** Box-plots represent coefficient of variation for each predicted posterior distribution in all test datasets. Colorbar represents variances of CHANGE-seq, GUIDE-seq, HEK293T, K562 and SITE-seq datasets.

### Effects of mismatches and off-target activity on the uncertainty of the predictions

To investigate whether crispAI-predicted off-target activity distributions captured effects of number of mis-matched positions in sgRNA-target pair and the ground truth assay detected off-target activity values for the CRISPR/Cas9 system, we used the left-out test portion of CHANGE-seq dataset and stratified the samples with respect to: (i) the base-pair mismatch count between the sgRNA and target sequences; and (ii) detected CHANGE-seq frequency and plotted the stratified folds with respect to the coefficient of variation of the associated crispAI-predicted off-target activity distributions.

We observed a consistent decrease in the coefficient of variation span as the mismatch count decreased from 6 to 0. (Fig. 5**b**). This result is expected since, as the number of allowed mismatches between sgRNA and target sequences increases, allowed degree of freedom for other sequence-based factors, that are shown to be correlated with higher off-target cleavage activity, increases (e.g., GC content, mismatch location) yielding a higher coefficient of variation in predicted off-target activity distributions.

**Fig. 5.**
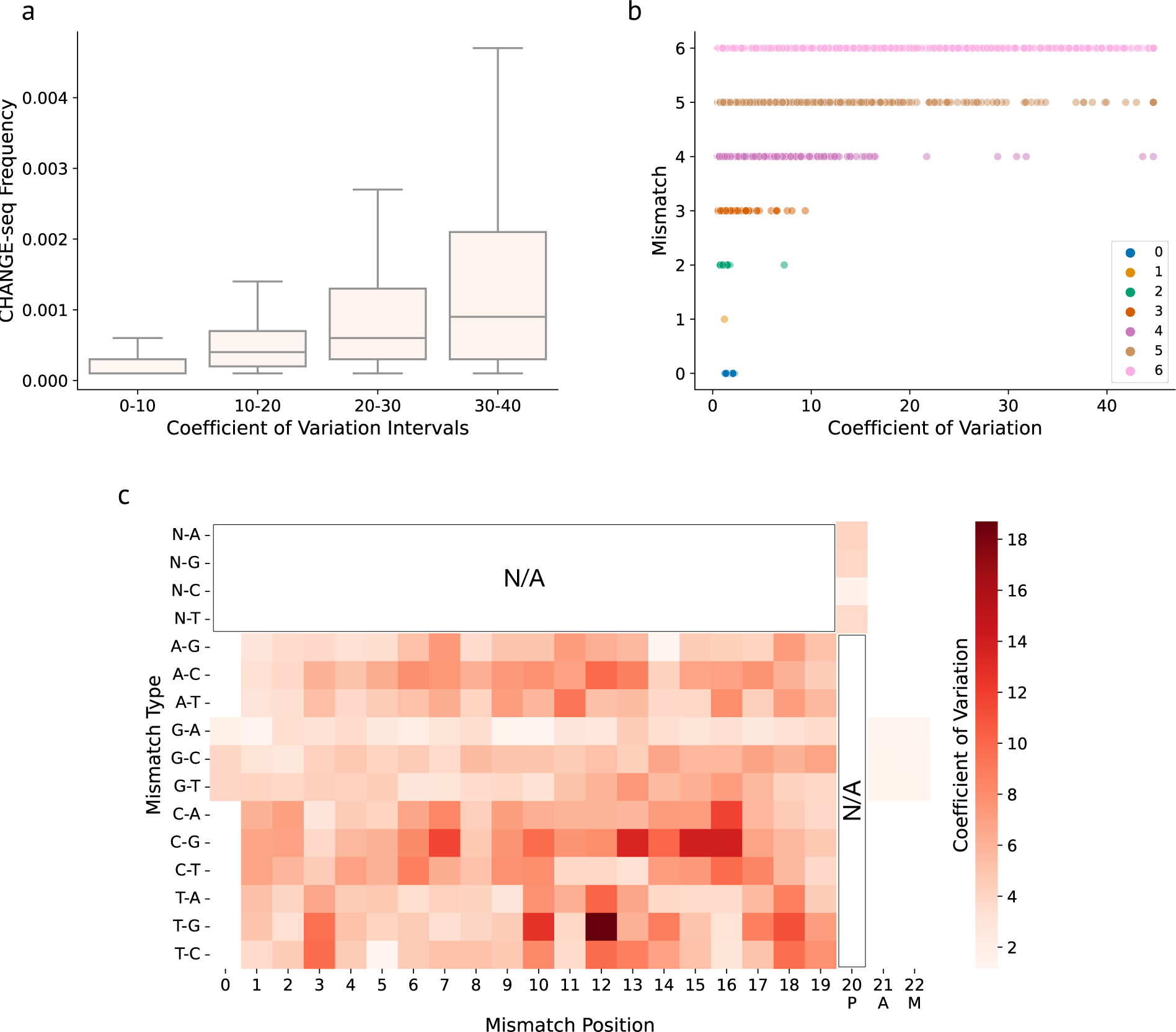
Similar to Fig. 4**d** coefficient of variation for each crispAI-predicted posterior off-target activity distribution on the test portion of the CHANGE-seq dataset are plotted against ground truth CHANGE-seq detected off-target activity and number of mismatched positions between sgRNA and DNA sequences of the associated sample. **a.** CHANGE-seq detected off-target activity values are depicted with box-plots for associated coefficient of variation intervals. Box-plots show the increase in CHANGE-seq detected off-target activity is associated with increase in coefficient of variation. **b.** Scatter plot illustrating the relationship between coefficient variation and number of mismatched positions between sgRNA-target pair indicating higher uncertainty as the number of mismatches increase. **c.** Grid-plot shows the coefficient of variation of the crispAI-predicted off-target cleavage activity distributions stratified with respect to types and positions of mismatches between sgRNA-target pairs on the test portion of the CHANGE-seq dataset. Certain mismatch types, such as T*→*G, and PAM-proximal mismatches yielded higher uncertainty.

We observed a similar trend for the ground truth CHANGE-seq detected activity values for 4 different coefficient of variation intervals (i.e., [0, 10], [10, 20], [20, 30], [30, 40]), where the detected frequency values for crispAI-predicted distributions with lower coefficient of variation values are higher than those with higher coefficient of variation values. (Fig. 5**b**) The relationship between ground truth CHANGE-seq detected off-target cleavage activity and the coefficient of variation of the crispAI-predicted distributions implies that as off-target cleavage activity decreases, our model’s confidence in its predictions for the associated sgRNA-target pair increases. This phenomenon is advantageous as it indicates that crispAI is more certain and consistent in identifying sgRNA-target pairs with lower off-target cleavage activity. Such sgRNA-target pairs exhibit more densely distributed predicted probability distributions, suggesting a greater level of confidence in the predictions.

To investigate the effcet of location and type of the mismatches on the uncertainty of the off-target cleavage activity, we stratified crispAI-predicted distributions on the test portion of the CHANGE-seq dataset with respect to: the type of the mismatch (e.g., A*→*C - sgRNA base is A and target base is C) and the position of the mismatch between sgRNA and target DNA sequences. Then we plotted the coefficient of variation of the crispAI-predicted distributions for the stratified folds in Fig.5. We observed an increase in coefficient of variation for PAM-proximal base loci as opposed to PAM-distal region. Additionally, we observed a significant increase in uncertainty for some mismatch types compared to others. Specifically: T *→* G, T *→* C and C *→* G sgRNA to target mismatch types. Recent studies widely reported that PAM-proximal mismatches are less tolerated for the cleavage activity [6, 33, 34] meaning that PAM-proximal sequence-based variations have higher effect. PAM-proximal mismatches yielded higher uncertainty in the crispAI-predicted cleavage activity distributions.

### crispAI-aggregate score enables uncertainty aware genome-wide sgRNA specificity prediction

We developed crispAI-aggregate, the first-of-its-kind genome-wide sgRNA specificity score, to provide aggregate score distributions for the sgRNA of interest. To calculate crispAI-aggregate distributions, we use Cas-OFFinder [35] to search for potential off-target sites of the sgRNA of interest up-to *N* mismatches, where *N* is a hyperparameter of the crispAI-aggregate score. Then the element-wise summation of crispAI-predicted posterior cleavage activity distributions for all obtained off-target sites are element-wise divided by the crispAI-predicted posterior cleavage activity distribution for the perfect homology target-site sequence (i.e., 0-mismatch target). Finally, crispAI-aggregate is defined as the logarithm of the obtained conditional distribution (Methods).

To evaluate crispAI-aggregate score, firstly we obtained the sgRNA specificity data curated by Fu et al. [19] on CHANGE-seq, TTISS [36] and GUIDE-seq datasets - providing specificity scores for 108, 59 and 10 sgRNAs respectively. We used sgRNAs in these dataset as inputs to crispAI-aggregate pipeline with maximum number of mismatches up-to *N* = 5 and obtained crispAI-aggregate distributions for all sgRNAs. To compare the performance of crispAI-aggregate score with other competing aggregation methods, we take expectations of the predicted distributions and obtained point predictions for the specificity aggregation task. Bar-plots in Fig.6**a** illustrates the performance of competing aggregate scores: MOFF-aggregate, CRISPR-Net, CFD, Elevation-aggregate, CRISPRoff, CNN std and crispAI-aggregate against Spearman correlation coefficient with respect to ground truth specificity values given in the curated dataset calculated based-on ground truth sequencing reads from respective *in vitro* assays. We observed that crispAI-aggregate significantly out-performed existing scores on CHANGE-seq and GUIDE-seq datasets with 14.27% and 6.99% improvements over best performing tools, MOFF and CRISPR-Net respectively and performed above average with 25.39% deterioration below MOFF-aggregate in Spearman correlation on sgRNAs in TTISS dataset, where average deterioriation among other methods is 41.26% below MOFF-aggregate score in Spearman correlation on this dataset.

Additionally, we visualised the Cumulative Distribution Functions (CDF) of the obtained crispAI-aggregate distributions for all 108 sgRNAs in the CHANGE-seq dataset in Fig.6**b** and annotated the CDFs with a colorbar depending on the associated ground truth specificity value. We observed that sgRNAs exhibiting higher specificity yielded crispAI-aggregate distributions with CDFs where the distribution is more densely populated around lower crispAI-aggregate score values (right-skewed PMFs), whereas sgRNAs exhibiting lower specificity yielded crispAI-aggregate distributions with CDFs where the distribution is more densely populated around higher crispAI-aggregate score values (left-skewed PMFs). This result supports our findings since lower crispAI-aggregate score values indicate higher specificity in sgRNAs.

To further evaluate the crispAI-aggregate score, we obtained all 2, 408 sgRNAs presented in the Avana library [12] targeting non-essential genes. Similarly to Fig.6, we obtained crispAI-aggregate distributions for all sgRNAs, again using *N* = 5. Then, we created 8 bins in total using the ground-truth Log Fold Change (LFC) values present in the Avana library obtaining bins from *−*2.3 to 0.9 LFC values associated with all analysed sgRNAs. The ridgeline plot in Fig.7**a** illustrates the expected values of obtained crispAI-aggregate distributions for each bin. Similarly to Fig.6**b** for sgRNAs with higher LFC values expected values of crispAI-aggregate distributions are more skewed and more densely populated around lower crispAI-aggregate score values. More specifically for the obtained bins we observed 6.092, 5.166, 4.454, 4.069, 3.373, 2.821, 2.840 and 3.333 mean crispAI-aggregate scores with *−*2.14, *−*1.66, *−*1.28, *−*0.89, *−*0.48, *−*0.09, 0.19 and 0.5 average LFC values respectively. Additionally, we measured MOFF-aggregate score for the same bins, bar plot in Fig.7**b** shows the average MOFF-aggregate scores 4.148, 3.575, 2.880, 2.266, 1.203, 0.234, 0.076 and 0.958 for each LFC bin in the respective order.

**Fig. 6.**
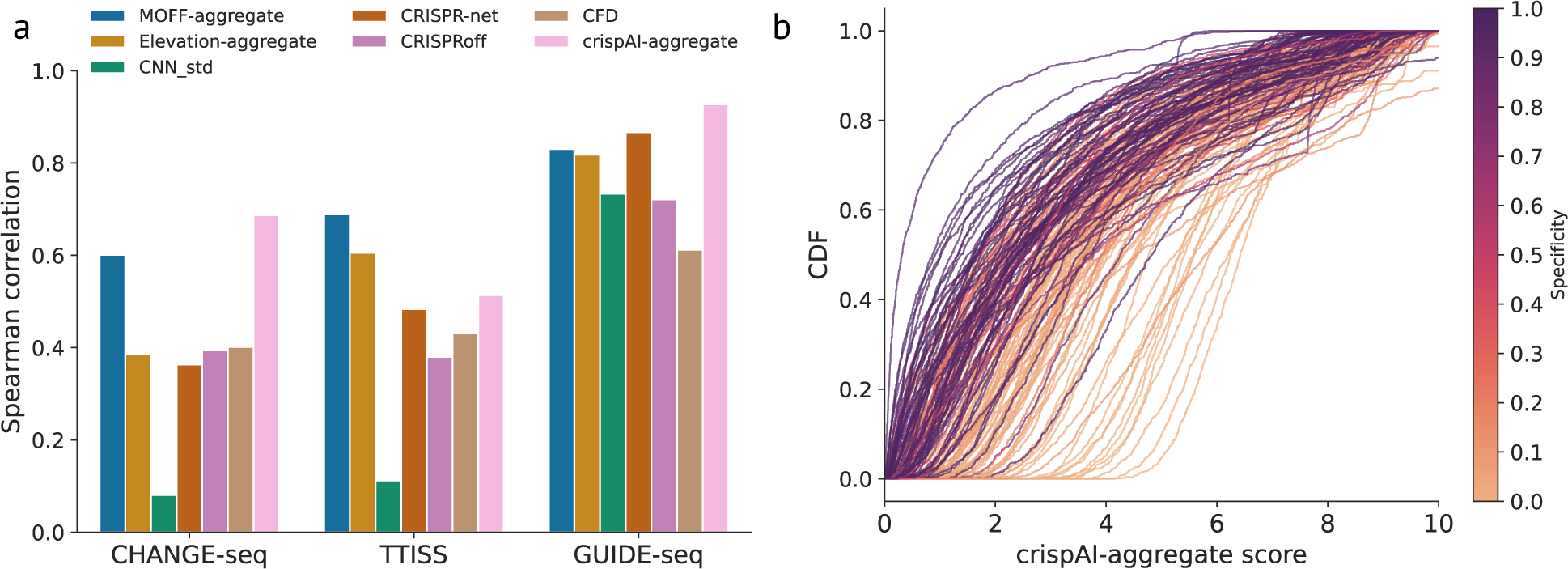
Genome-wide uncertainty aware sgRNA specificity prediction with crispAI-aggregate score. **a.** Genome-wide sgRNA specificity score, crispAI-aggregate score, is defined as the logarithm of the ratio between sum of crispAI-predicted off-target scores up-to *N* mismatches and the on-target sequence, where *N* is a hyperparameter of the score. For the plots herein *N* = 5 is used. **a.** Bar-plot represents the Spearman correlation between sgRNA specificity, as presented in [19], and predicted aggregate scores by MOFF-aggregate, CRISPR-Net, CFD, Elevation-aggregate, CRISPRoff, CNN std and crispAI-aggregate on CHANGE-seq, TTISS and GUIDE-seq datasets with *n* = 108, 59 and 10 sgRNAs respectively. **b.** Cumulative Distribution Functions (CDFs) of predicted crispAI-aggregate score distributions are depicted. The colorbar represents the sgRNA specificty of the associated CDF in the respective dataset.

The genome-wide specificity distributions produced by crispAI-aggregate enables distinguishing between sgRNAs with similar point predictions. To demonstrate this histograms in Fig.7**c** depicts crispAI-aggregate score distributions for two sgRNAs targeting CSH1 and SPACA7 genes with LFC values *−*0.305 and *−*0.294 in the Avana library and the mean values of crispAI-aggregate scores are calculated as 4.055 and 4.012 respectively. Although both the associated LFC values and the expected values of the predicted crispAI-aggregate distributions are very close for these sgRNAs, obtained distributions are different. Specifically, the sgRNA targeting CSH1 gene yielded a wider crispAI-aggregate score distribution compared to the sgRNA targeting SPACA7 gene with coefficient of variation values of 0.804 and 0.334. This finding suggests crispAI-aggregate score enables prioritization among sgRNAs with similar point predictions by providing richer information for genome-wide specificity prediction problem.

**Fig. 7.**
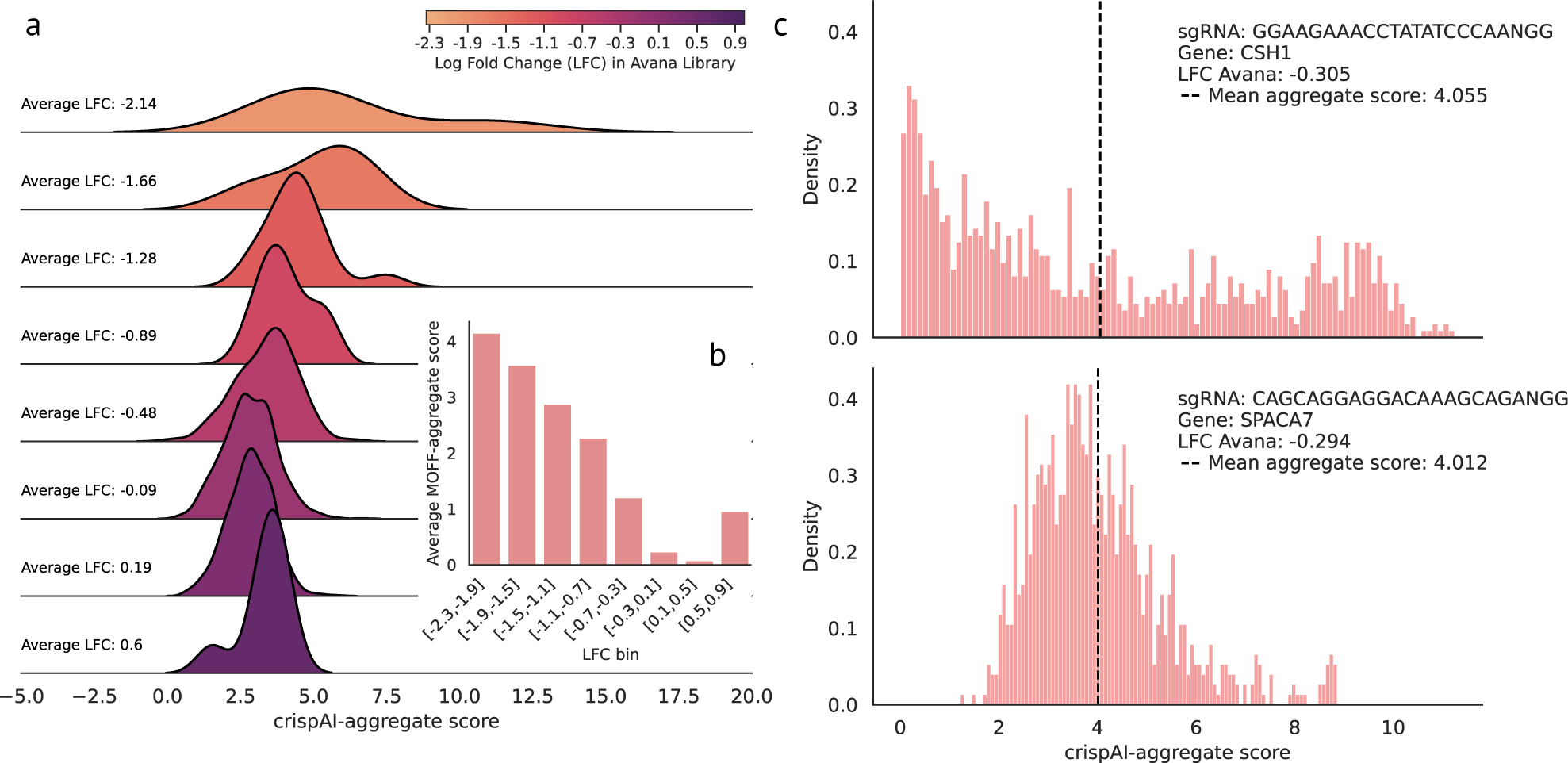
Prioritization of sgRNA specificity with crispAI-aggregate score. **a.** Ridgeline plot for expected values of crispAI-aggregate score distributions on 2, 408 sgRNAs given in the Avana library [12]. First, we searched genome-wide for up to 5 mismatch off-target sites for all 2, 408 sgRNAs using CasOffFinder, obtaining *n* samples in total, and calculated crispAI-aggregate scores for each sgRNA. Then, we used associated Log Fold Change (LFC) values for each sgRNA to obtain 8 bins with different bin ranges based on LFC values. The ridgeline plot depicts expected crispAI-aggregate score for each LFC bin. **b.** Bar-plot representing the average MOFF-aggregate scores for the bins used in **a**. **c.** Histograms depict crispAI-aggregate score for two similar LFC sgRNAs targeting CSH1 and SPACA7 genes, with *−*0.305 and *−*0.294 LFC values in the Avana Library respectively. The expected value of predicted crispAI-aggregate score distributions are 4.055 and 4.012 in the respective order.

## Discussion

The development of *in silico* predictive models for CRISPR/Cas9 off-target activity prediction has achieved significant milestones in various approaches, including heuristic models, traditional machine learning models and deep learning models. Heuristic models were among the early attempts to predict off-target activity [12,29,37,38], relying on predefined rules and sequence patterns. While these models provided initial insights, they often lacked generalizability and accuracy. The advent of learning models, such as traditional machine learning algorithms like SVM, random forest, and logistic regression, brought improvements by leveraging data-driven approaches [14,39–41]. These models incorporated features derived from sequence characteristics and demonstrated enhanced prediction capabilities. However, with the emergence of deep learning models in the field, including convolutional neural networks (CNNs) and recurrent neural networks (RNNs), the effectiveness and prediction capabilities of the modelling efforts significantly improved [16, 26, 32, 42]. Deep learning models could effectively handle large volumes of complex data, capturing intricate patterns and achieving superior performance in predicting off-target activity. The development of deep learning-based prediction models also enabled utilization of physical features modelling the off-target activity problem with more depth [17]. Accurately quantifying uncertainty in off-target activity predictions is a crucial next milestone this study aims to address. While predictive models have shown promising results in identifying potential off-target cleavage activity, they provide point predictions without considering the associated uncertainty. Incorporating uncertainty estimates into these models would provide a more comprehensive understanding of the reliability and confidence of the predictions. It would enable researchers and practitioners to differentiate between sites with similar point predictions but different levels of uncertainty, allowing for better risk assessment and prioritization of potential off-target sites. Additionally, quantifying uncertainty would enhance the transparency and communication of the prediction results, providing stakeholders with a clearer understanding of the associated risks.

For sequencing data analysis, it is essential to acknowledge the heterogeneity of zeros, as they can originate from diverse processes, introducing noise and uncertainty into the data. Thorough modeling is required to address this phenomenon and ensure accurate interpretation of results. Two primary categories of zeros are encountered: technical zeros and biological zeros [23]. Technical zeros arise from limitations in sample preparation or sequencing, leading to partial or complete reduction in countable sequences. Examples include biases in amplification, sequencing depth limitations, or batch effects. In contrast, biological zeros occur when a specific sequence is genuinely absent from the biological system under investigation, such as unique bacterial strains in different individuals or gene deletions in knockout experiments. Extensive research efforts have been devoted to overcoming this type of challenge in various application domains. Researchers have developed sophisticated techniques, including the use of technical components like zero-inflated distributions, to effectively model noisy zero counts. These approaches have been applied in domains such as microbiome [43] studies, single-cell RNA sequencing [24, 44], and bulk RNA sequencing [45], enabling improved accuracy and insight in differential gene expression analysis [46] and facilitating a deeper understanding of biological processes. In off-target activity detection data, we use technical zeros to refer to the off-target sites that can not be captured due to the limited sensitivity of the detection assays. To address the technical zero problem in the raw count version of off-target activity data, we employed a count noise-modeling approach that utilized a Zero-Inflated Negative Binomial (ZINB) distribution. This allowed us to accurately model the characteristics of the data, account for excessive zeros, and incorporate uncertainty modeling.

Although our approach provides a more comprehensive risk assessment than traditional point predictions, it is important to note that our modelling effort tackles the uncertainty problem by considering abundant number of potential off-target sites, which is not suitable for modelling uncertainty in the on-target cleavage activity. Hence, a limitation of crispAI is that it is not designed for on-target activity prediction. The current model does not share the limitation of all sequence-based models, whose predictions are solely based on the sgRNA-target DNA sequence pair because they cannot differentiate between off-target activities of identical sgRNA-target interfaces at different genomic loci. Whereas, crispAI incorporates physical features of the genomic loci at the target region, using richer information compared to sequence-based models and allowing differentiation between target regions with the same off-target sequence. Therefore, our method differentiates from the other uncertainty aware Gaussian Process Regression (GPR)-based method [22] by sampling uncertainty estimates using physical descriptors in addition to sequence-based features of the sgRNA-target context whereas GPR-based model is unable to distinguish between target sites with the same off-target sequence.

We observed an increase in the coefficient of variations for the predicted activity distribution of sgRNA-target DNA interfaces that had mismatches in the PAM-proximal region compared to the coefficient of variation we observed for predicted activity distribution for interfaces with PAM-distal mismatches. This observed difference in the coefficient of variation for the predicted distributions for the said interfaces is in concordance with the different roles of PAM-proximal and PAM-distal base pairings have in the mechanism of the CRISPR/Cas9 based editing. Correct PAM-proximal base pairing is essential for initiating sgRNA-target DNA heteroduplex formation and therefore the stable binding of the CRISPR/Cas9 complex to the target DNA loci. Therefore PAM-proximal mismatches can adversely affect correct binding and subsequently off-target activity at the given loci [47]. At the same time initiation of PAM-proximal base pairing is a stochastic process and this may explain why it is more challenging to correctly predict the effect of PAM-proximal mismatches with high certainty [48].

## Material & Methods

### Datasets

For the training of crispAI, we used CHANGE-seq dataset [9]. CHANGE-seq is a scalable, automatable tagmentation-based method for measuring the genome-wide activity of Cas9 *in vitro*. Authors identified 201, 9434 off-taget sites on 110 sgRNAs across 13 therapeutically relevant loci in human primary T cells. Although CHANGE-seq assay is a high sensitivity *in vitro* genome-wide off-target detection method [20], it is still not able to detect all of the potential off-target sites due to limited sensitivity of the experimental apparatus. However, studies suggested that off-target sites which have several mismatched positions with the respective sgRNA sequence (i.e., up-to 6 base-pairs) are *putative* off-target sites and are potentially harmful.

Many genome alignment-based methods [35, 38, 49] have been proposed for *in silico* discovery of the putative off-target sites for a given sgRNA. We employed one of the most popular, light-weight tool CasOFFinder [35] for this task owing to its ease of use and search speed. Specifically, we searched for putative off-target sites for all 110 sgRNAs presented in CHANGE-seq dataset with up to 6 allowed base-pair mismatched positions with the sgRNA sequence. We obtained a total of 1, 783, 801 putative off-target sites, yielding a total of 1, 581, 757 off-target sites that are not also in CHANGE-seq dataset.

To evaluate crispAI, we used 5 different test sets obtained with different assays. More specifically: (i) we randomly splitted 10% of the CHANGE-seq dataset on human primary T-cells obtaining 168, 465 off-target sites for *n* sgRNAs; (ii) GUIDE-seq dataset containing 443 off-target sites for *n* sgRNAs; (iii) SITE-seq dataset containing 6, 097 off-target sites for *n* sgRNAs; (iv) HEK293T and (v) K562 cell-lines datasets used in Chuai et al. [32] containing 536 and 120 off-target sites for 12 and 18 sgRNAs respectively.

### Modelling of crispAI

#### Problem formulation

Letting *x_s_ ∈ {*0, 1*}^ms×ℓ^*, where *m_s_* and *ℓ* are feature dimension and sequence length respectively, be the nucleotide sequence-based features of sgRNA-target pair interface, *x_p_ ∈* [0, 1]*^mp×ℓ^*, where *m_p_* is feature dimension for physical features, denote the physical descriptor based features of sgRNA-target pair interface and *y ∈* R denote the cleavage read depth, we model off-target activity data generation process with a Zero-Inflated Negative Binomial (ZINB) distribution. The ZINB distribution is an appropriate choice for modeling count data that is both highly sparse and overdispersed. A ZINB mixture model can be constructed using two components: a point mass at zero, which represents the excessive number of undetected inactive samples in the data, and a negative binomial component that models the count distribution. In the context of off-target assay data for CRISPR-based editing technologies, the point mass at zero is expected to capture the abundance of undetected inactive samples, while the negative binomial component is used to represent the sequencing reads for active samples. Hence, we model *y* with the following set of parametric equations as:

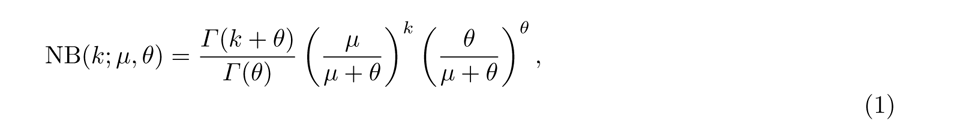

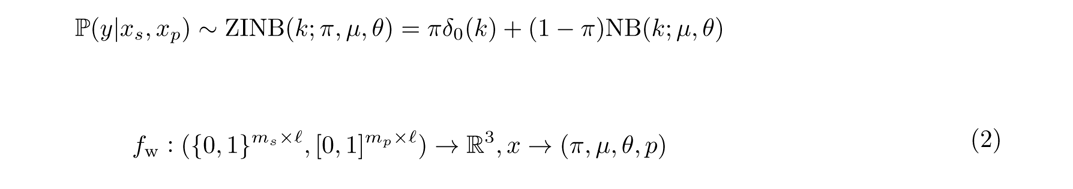

where ZINB(*k*; *π, µ, θ*) is the ZINB distribution with parameters *π*, *µ* and *θ* for the mixture coefficient representing the point-mass (*δ*_0_) at 0, mean and dispersion of the negative binomial component (NB), respectively. We model the parameters for the conditional distribution with a multi-input, multi-output parametric function *f*_w_ : (*{*0, 1*}^m×ℓ^,* [0, 1]*^mp×ℓ^*) *→* R^3^ with parameter set w. Note that the output space of the function *f*_w_ is R^3^, where three output dimensions represent *π*, *µ* and *θ*. Thus, using a data-set consisting of *N* samples in the form of 3-tuples, *D* = *{*(*x^i^, x^i^, y^i^*)*}^N^*, we want to be able to calculate the posterior distribution of the cleavage score *y* given the features *x_s_*, *x_p_* and the data *D* — that is P(*y|x_s_, x_p_, D*).

#### Encoding of the sequence-based sgRNA-target interface features

We used 4-bit one-hot-encoding vectors representing the letters in the alphabet *{A, G, C, T}*. Using the one-hot-encoded representations for each base in sgRNA and target sequences yields 4 *×* 23 binary matrices for each sequence. Then we employed the sgRNA-target pair encoding scheme proposed in Lin et al. [26]. We represent any sgRNA-target sequence pair with a 6 *×* 23 matrix as follows: First, both sgRNA and the target sequence is one-hot encoded. Then, obtained binary matrices are merged via an element-wise OR operation. Hence, resulting 4 *×* 23 binary matrix shows the mismatches between sgRNA-target sequence pair. However, OR operation does not preserve the direction of the mismatch. To help ameliorate this information loss, a two-bit direction channel is concatenated to the resulting binary matrix. For example, at a base-pair loci, ‘0011 *−* 10’ represents the mismatch ‘*G → C*’; ‘0011 *−* 01’ represents the mismatch ‘*C → G*’ and one-hot vector ‘0100 *−* 00’ represents the matched loci ‘*T → T* ’, obtaining a 6 *×* 23 matrix for the sequence-based features of the sgRNA-target pairs.

#### Physical descriptors and encoding of the 147-bp sequence context of the target site

We used 147-bp sequence context on the off-target loci (i.e., 73-bp flank on each side of an off-target sequence position) with a sliding window approach to obtain: (i) Nucleotide BDM score [27]; (ii) GC content; (iii) NuPoP occupancy and (iv) NuPoP affinity scores [28] and normalized the obtained scores in the [0, 1] interval. Hence, we obtained 4 *×* 23 matrix for physical descriptors of the off-target sequence context. GC count refers to the proportion of G and C bases within the 147-bp sliding window, centered around a given off-target sequence position. Nucleotide BDM is a training-free method to approximate the algorithmic complexity of a given DNA sequence. NuPoP scores refer to a Hidden Markov Model (HMM), trained to estimate the nucleosome affinity and occupancy at single base-pair resolution. For a more detailed discussion on the effects of these features to CRISPR-based cleavage activity, we refer to Störtz et al. [17].

#### Architecture and training of crispAI

crispAI is an end-to-end multi-output CNN and bi-LSTM fusion neural network specifically designed to quantify the uncertainty in off-target cleavage activity of sgRNA-target pairs for CRISPR/Cas9 system. We show that crispAI architecture is able to increase performance on off-target cleavage activity prediction task on different test sets, while providing uncertainty quantification in off-target activity (Results).

crispAI architecture is designed to estimate the parameters of the ZINB distribution conditioned on the sgRNA-target interface features. First, we input binary matrix encoding of the interface to a series of Convolutional Neural Network (CNN) and bi-directional Long-Short Term Memory (bi-LSTM) layers simultaneously for both spatial and temporal feature extraction. Specifically, we use 2 consecutive CNN layers with 128 and 32 kernels with 1 and 3 filter sizes respectively and a bi-directional LSTM layer with 128 hidden neurons in each direction. The output activations of the CNN layers are batch-normalized while the output of the last CNN layer is pooled with a MaxPool layer of kernel size 2 and with a stride of 2. The obtained encodings are further processed with two distinct 128-neuron Fully Connected (FC) layers individually. We then concatenate the final encodings outputted by the FC layers into a single 256-dimensional vector. The final FC layer with 64 neurons processes the concatenated vector and is inputted to 3 output neurons, one for each parameter of the ZINB distribution.

We used ReLU activations for FC and CNN layers. The activation function choices for the individual output neurons predicting 3 associated parameters for the ZINB distribution, *π*, *µ* and *θ*, are discussed below. The architecture can be formulated through following equations:

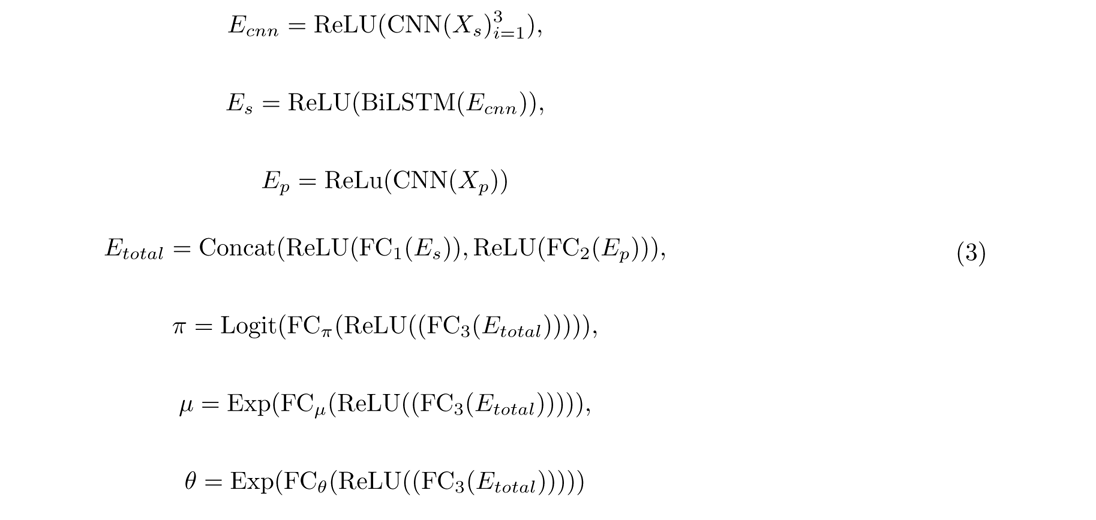

where *E_cnn_, E_s_, E_p_* and *E_total_* represent CNN encoding of the sequence-based features, biLSTM encoding of the CNN extracted features, CNN encoding of physical descriptors and concatenated total encoding of the sequence-based and physical descriptor features respectively. To ensure positivity on parameters *π* and *µ*, we use exponential activations, and for drop-out parameter *π*, we use Logit activation for ease of integration with Negative Log Likelihood (NLL) loss for ZINB distribution.

All network parameters in (1), are learned with a multi-task training framework using Stochastic Gradient Descent (SGD) algorithm. To train the network weights, we use Negative Log Likelihood Loss of ZINB (*L*_ZINB_ = NLL_ZINB_) for three parameters of the ZINB distribution.

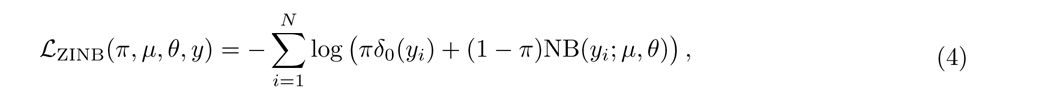

where all parameters in (4) are defined in Problem Formulation section.

We trained the crispAI architecture with the same training settings on all datasets. We splitted CHANGE-seq and GUIDE-seq dataset into training, validation and testing sets with 70%, 20% and 10% respectively. For

DeepCRISPR data we did not use a validation set and used the same hyper-parameter configurations we discovered using the CHANGE-seq validation set. Hence, we applied a 80%-20% train-test split on the latter.

We used Adam optimizer [50] for optimization with a learning rate of 0.00001. We stopped the training with early stopping, watching either the validation or training loss for 50-epochs, with a maximum epoch number of 500. We implemented crispAI on Python 3.9 using PyTorch [51]. Finally, we used SCVI-tools [52] to implement the loss functions. The model is trained on a single 24G NVIDIA TITAN RTX GPU.

#### Genome-wide sgRNA specificity prediction with crispAI-aggregate score

We designed, the first of its kind, crispAI-aggregate score for uncertainty-aware sgRNA genome wide specificity prediction, inspired by

MOFF-aggregate score [19]. First, we use CasOFFinder [35] to search putative off-target DNA sites up-to a user specified (i.e., *N* = 5) number of mismatches genome-wide for the sgRNA of interest. Then, we obtain crispAI-predicted cleavage activity distributions for each obtained putative off-target site. Then, we calculate the ratio between the summation of the crispAI-predicted distributions of all of the detected off-target sites and the crispAI-predicted distribution for the on-target site. Finally, crispAI-aggregate score for a given sgRNA is defined as the log of the obtained ratio.

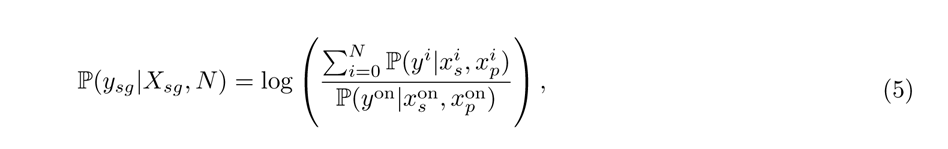

where *y_sg_, X_sg_* and *N* denote genome-wide specificity score, sequence-based features of the sgRNA of interest and user specified maximum number of mismatches to search for genome-wide (5 by default). The variables on the right hand side of (5) are defined in (1) and indexed by the off-target site indice in the nominator and the on-target site indice in the denominator.

## Code and Data Availability

All necessary scripts and data to replicate the results presented in figures are deposited to Zenodo https://zenodo.org/records/10516069?token=eyJhbGciOiJIUzUxMiJ9.eyJpZCI6ImNmZjIxNWE0LTFmMTUtNGM5ZC1hYTliLWRiMmIwZRNooX9OxDtMxVqnyuc78HX43FSY5pFY5vvY4dD5jD-Ib8IbS1VfUYDMhj6jnYA-jdCO21-IX4m6tkp7_ihpFCQ, the tool and the trained model are available at https://github.com/furkanozdenn/crispr-offtarget-uncertainty.

## Competing Interests

Authors declare no competing interest.

## Author Contributions

FO and PM designed the study. FO designed and implemented the model and performed the experiments. FO and PM wrote the manuscript.

## Author Information

Peter Minary is a Research Lecturer at the Department of Computer Science, University of Oxford. His research interests include computational (structural) biology, machine learning and CRISPR-based genome editing technologies.

Furkan Ozden is a DPhil student in Computer Science at the University of Oxford. His research interests include machine learning, genomic variation and CRISPR-based genome editing technologies.

## Rights Retention Statement

FO acknowledges the funding from Google DeepMind. For the purpose of Open Access, the author has applied a CC BY public copyright licence to any Author Accepted Manuscript (AAM) version arising from this submission.

## Notes

### Competing Interest Statement

The authors have declared no competing interest.

### Summary of Updates

- Added physical descriptors and changed neural network architecture. - Proposed uncertainty aware genome-wide sgRNA specificity score - Added additional performance experiments

